# Structural and functional characterization of the KHNYN extended-diKH domain for mediating ZAP antiviral activity

**DOI:** 10.1101/2024.12.23.630104

**Authors:** Rebecca L Youle, María José Lista, Clement Bouton, Simone Kunzelmann, Harry Wilson, Matthew A Cottee, Andrew G Purkiss, Elizabeth R Morris, Stuart J D Neil, Ian A Taylor, Chad M Swanson

## Abstract

Zinc finger antiviral protein (ZAP) binds CpG dinucleotides in viral RNA and targets them for decay. ZAP interacts with several cofactors to form the ZAP antiviral system, including KHNYN, a multidomain endoribonuclease required for ZAP-mediated RNA decay. However, it is unclear how the individual domains in KHNYN contribute to its activity. Here, we demonstrate that the KHNYN amino terminal extended-diKH (ex-diKH) domain is required for antiviral activity and present its crystal structure. The structure belongs to a rare group of KH-containing domains, characterized by a non-canonical arrangement between two type-1 KH modules, with an additional helical bundle. N4BP1 is a KHNYN paralog with an ex-diKH domain that functionally complements the KHNYN ex-diKH domain. Interestingly, the ex-diKH domain structure is present in N4BP1-like proteins in lancelets, which are basal chordates, indicating that it is evolutionarily ancient. While many KH domains demonstrate RNA binding activity, biolayer interferometry and electrophoretic mobility shift assays indicate that the KHNYN ex-diKH domain does not bind RNA. Furthermore, residues required for canonical KH domains to bind RNA are not required for KHNYN antiviral activity. By contrast, an inter-KH domain cleft in KHNYN is a potential protein-protein interaction site and mutations that eliminate arginine salt bridges at the edge of this cleft decrease KHNYN antiviral activity. This suggests that this domain could be a binding site for an unknown KHNYN cofactor.

## Introduction

ZAP (also called ZC3HAV1 or PARP13) is an RNA binding protein with antiviral activity against diverse virus families (1–4). While ZAP has multiple mechanisms by which it restricts viral replication, the best characterized are translation inhibition and RNA decay (1, 3–5). The ZAP RNA binding domain (RBD) binds CpG dinucleotides with high affinity (6–8). When spaced approximately 15-30 nucleotides apart in an A/U-rich context, clusters of more than 15 CpGs constitute a ZAP-response element (ZRE) in viral RNA (6, 9). ZAP evolved in tetrapods and is part of a core set of evolutionarily conserved interferon-stimulated genes (ISGs) (10, 11). Many pathogenic human viruses have evolved to evade its activity by encoding protein countermeasures or suppressing CpG abundance in their genomes (3, 4, 6). Synonymous mutations that create CpGs impose a fitness cost for diverse viruses, including HIV-1 (12, 13). CpG abundance in HIV-1 is particularly low, likely due to evolutionary pressure from ZAP antiviral activity, and its abundance in HIV-1 *env* is linked to disease progression (14–17).

KH and NYN containing protein (KHNYN) was recently shown to be a co-factor for ZAP-directed inhibition of retroviral replication (18). KHNYN interacts with ZAP in an RNA-independent manner and KHNYN depletion increases retroviral gene expression and infectious virus production for both murine leukemia virus and HIV-1 that has CpGs introduced via synonymous genome recoding to create a ZRE (18, 19). Additionally, KHNYN displays antiviral activity against ZAP-sensitive strains of human cytomegalovirus (20). However, as KHNYN has only recently been identified as a ZAP cofactor and has no other identified function, it remains poorly characterized. KHNYN contains three structural domains, an N-terminal KH domain, a PilT N-terminus (PIN) nuclease domain, and a C-terminal CUE_like_ domain (also known as a CUBAN domain) that binds ubiquitin and NEDD8 (18, 21–23). The CUE_like_ domain also contains a C-terminal nuclear export signal required for KHNYN cytoplasmic localization, its interaction with ZAP and antiviral activity (24). KHNYN has two human paralogs, NYNRIN and N4BP1 (25). NYNRIN has been proposed to regulate trophoblast invasion during placental development while N4BP1 inhibits viral replication and regulates innate immune signaling (26–32).

We have previously shown that the catalytic residues in the KHNYN PIN domain are required for its antiviral activity and the CUE_like_ domain regulates both KHNYN abundance and subcellular localization (18, 24). However, the role of the N-terminal KH domain remains unclear. Initially described as an unconventional KH domain (21), this domain is present only in KHNYN, N4BP1 and NYNRIN orthologs (33). Our previous evolutionary and AlphaFold2 analysis of the KHNYN gene family indicated that the N-terminal KH domain is comprised of two tandem canonical type-1 KH domains (KH1 & KH2) each with a β-α-α-β core (34) with an additional extension C-terminal to the second KH domain (24).

To determine the functional requirement of the KHNYN KH domains in ZAP-directed antiviral activity, we carried out a combined virological and structural study. Deletion of the KH region resulted in loss of KHNYN antiviral activity. The crystal structure revealed an orthogonal arrangement of the KH1 and KH2 modules and extended C-terminal alpha helical bundle not observed in other tandem KH domains, giving rise to the term extended-diKH (ex-diKH) domain. The N4BP1 ex-diKH domain has a similar structure and can replace the KHNYN ex-diKH domain without abrogating antiviral activity. While most KH domains bind short nucleic acid sequences, the GxxG motif that makes core RNA-protein interactions is absent from KHNYN KH1 and our binding studies showed no evidence of RNA binding activity by the ex-diKH. However, mutations in arginine residues that support an interdomain KH1-KH2 cleft resulted in loss of antiviral activity. Therefore, we hypothesize that the KHNYN ex-diKH domain is a protein-protein interaction site required for KHNYN antiviral function.

## Results

### The KHNYN extended-diKH domain is required for antiviral activity

KHNYN was first annotated in the SUPERFAMILY database to contain a single KH domain (residues 58-141). Deletion of this region decreased antiviral activity against CpG-enriched HIV (HIV-CpG). However, it localized the protein to cytoplasmic puncta not present for wild-type KHNYN (18), suggesting that this deletion negatively impacted protein folding, resulting in aggregation. Furthermore, our subsequent Alphafold2 and evolutionary analysis predicted that the KHNYN N-terminal region actually contained a structured region of approximately 200 amino acids, which included two KH modules and a further C-terminal helical bundle comprising an extended-diKH domain (ex-diKH) (**Fig 1A**) (24, 35, 36). The structural homology-based tool Phyre2 (37) also predicted an ex-diKH domain similar to the AlphaFold2 model based on homology to a N4BP1 structure (PDB 6q3v) that shares 37% identity to the KHNYN domain.

**Figure 1.**
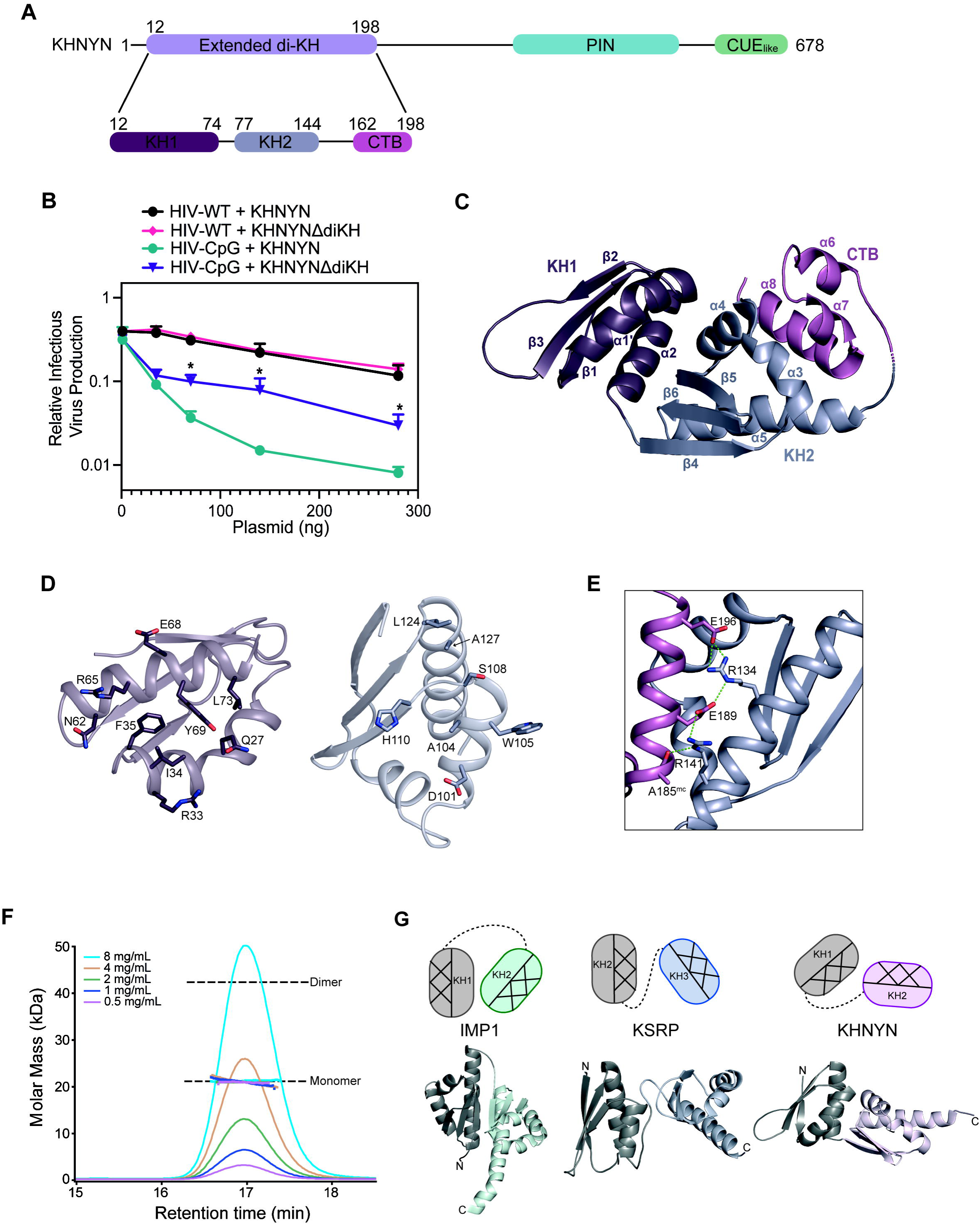
The KHNYN ex-diKH domain is required for antiviral activity and is a member of a novel group of diKH domain structures. (**A**) (upper) schematic representation of KHNYN. The positions of structural domains are highlighted; lilac (ex-diKH, residues 12-198), cyan (PIN) and green CUE_like_. (Lower) schematic of the ex-diKH domain, purple (KH1, residues 12-74), blue grey (KH2, residues 77-144) and magenta (CTB, residues 162-198) (**B**) Infectious virus production from KHNYN CRISPR HeLa cells co-transfected with pHIV-WT or pHIV-CpG and increasing amount of CRISPR-resistant flag-tagged pKHNYN or pKHNYNΔdiKH expressing plasmids. Each point shows the average value of three independent experiments normalized to the value obtained for HIV-WT in the absence of KHNYN. (**C**) Crystal structure of ex-diKH. The protein backbone is shown in cartoon, domains are colored as in **A**, secondary structure elements are labeled from N- to C-termini. (**D)** Details of the KH1-KH2 interface. Cartoon representations of the protein backbone of KH1 (left) and KH2 (right) are shown, colored as in **C**. The view is of from one domain looking into the interface of the other. Residues with side chains that make hydrogen bonding and hydrophobic interactions that contribute to interdomain packing (KH1: Q27, R33, I34, F35, N62, R65, E68, Y69, L73) and (KH2: D101, A104, W105, S108, H110, A127, L124) are shown as sticks, colored by atom type. (**E**) The KH2-CTB interdomain interface. KH2 and the CTB are shown in cartoon representation, colored as in **C**. Residues R134, A185, E189 and E196 that make hydrogen bonds, represented by the green dashed lines, are shown as sticks, mc = mainchain. (**F**) SEC-MALLS analysis of ex-diKH. The sample loading concentrations were 8 mg/mL (cyan), 4 mg/mL (wheat), 2mg/mL (green), 1 mg/mL (blue) and 0.5 mg/mL (magenta). The differential refractive index (dRI) is plotted against column retention time and the molar mass, determined at 1 s intervals throughout the elution of each peak, is plotted as points. The monomer and dimer molecular mass for ex-diKH are indicated by the dashed lines. (**G**) diKH domain arrangements. diKH domains from IMP1, KSRP and KHNYN are shown in cartoon colored grey & mint, grey & light blue and grey & pink respectively, N- and C-termini are labeled. The relative orientation of KH domains in each structure is shown above schematically, color coded to the corresponding domain. The helical and β-sheet faces of each KH domain are shown hatched and plain respectively.

To determine if the ex-diKH domain is required for KHNYN antiviral activity, we deleted it (KHNYNΔdiKH) and analyzed whether this mutant protein restricted ZAP-sensitive HIV-CpG, which contains a ZRE introduced into *env* through synonymous mutagenesis (6, 18). As a control, we also tested both wild-type KHNYN and KHNYNΔdiKH on wild-type HIV-1 (HIV-WT), which is ZAP-resistant due to low CpG abundance (6, 18). When KHNYN or KHNYNΔdiKH were expressed in KHNYN CRISPR cells, KHNYN potently inhibited HIV-CpG while KHNYNΔdiKH had reduced antiviral activity (**Fig 1B** and **Fig S1A**). This shows that the ex-diKH domain is required for full KHNYN antiviral activity. Importantly, we found that the ex-diKH domain was not required for the interaction of KHNYN with ZAP, KHNYN steady-state subcellular localization or trafficking to the nucleus (**Fig S1B-C**).

### Novel architecture of KHNYN extended-diKH domain

We then determined a crystal structure for the KHNYN ex-diKH domain (**Fig 1C, Fig S1D** and **Table S1**). The asymmetric unit (ASU) contains two copies of the ex-diKH domain (**Fig S1D**). In each copy, the domain spans residues 12-198 and is composed of an N-terminal type-1 KH module (KH1, residues 12-74), a short linker (residues 75 and 76), a second type-1 KH module (KH2, residues 77-144), a longer linker (residues 145-161) and a C-terminal bundle (CTB) extension composed of three alpha helices (residues 162-198) (**Fig 1C**). The KH2 domain has the expected β1-α1-α2-β2-β3-α3 type-1 KH topology but in KH1 α1 and α2 are near continuous constituting α1’ that contains a pronounced proline kink surrounding P29. Inspection of the domain packing within the protein revealed a substantial interdomain interface that packs KH1 onto KH2 (**Fig 1D**). This interaction is stabilized through multiple side chain interactions between residues on α1 and α2 of KH1 and α4, the α4-β5 linker and α5 of KH2. At the center of the interface, these include hydrophobic packing of KH1 side chains I34, F35, Y69 and L73 with A104, H110, L124 and A127 of KH2. These are accompanied by surrounding salt bridges and hydrogen bonding between R33 and D101, R65 and the mainchain carbonyl of S106 and between Y69 and H110, with R33 also π-stacking with W105. Additionally, the CTB extension packs onto the KH2 domain (**Fig 1E**) making an interface stabilized by multiple interactions between CTB α8 and KH2 α5, including a salt bridge network between E196 and R134, R134 and E189 and R141 as well as hydrogen bonding between R141 and the carbonyl of the mainchain A185.

While there were two copies of the ex-diKH in the ASU (**Fig S1D**), assessment of the protein-protein interface using PDB-PISA interface analysis (38) revealed only minimal contacts with a buried surface area of 393 Å^2^. The computed Complex Formation Significance Score of 0.0 also suggested that the interface is unlikely to have biological relevance. This conclusion is supported by solution SEC-MALLS data (**Fig 1F**) that revealed that the ex-diKH domain was exclusively monomeric over the range of concentrations tested (0.5 – 8 mg/mL). Therefore, the dimer observed in the crystal likely is a result of crystal packing rather than any residual assembly associated with an ex-diKH domain homo-oligomerization function.

A KHNYN ex-diKH structural similarity search using the DALI server (39, 40) identified the N4BP1 ex-diKH (PDB 6q3v) as the top overall match with an RMSD (Å) of 1.8 and a Z score of 23.9 over 174 Cα positions. In addition, the search also identified matches with single KH domains from a variety of RNA-binding proteins containing multiple KH domains with Z scores ranging from 8 to 9.6 (**Table S2**). Analysis of the arrangement and relative orientation of the two KH modules in the KHNYN ex-diKH structure in comparison to other diKH structures further allowed classification of three major groups of type-1 diKH arrangements (**Fig 1G**). The first and most common interdomain arrangement of type-1 diKH modules observed is the side-by-side anti-parallel conformation seen in IMP1 KH1-KH2, Nova-1 and PCBP2 (41–43). In this arrangement, the three β-strands of each KH module align to form an extended sheet made of six β-strands, with stabilization from interactions between the α3 helix in each KH module (34). The second group of interdomain arrangement is the orthogonal linker arrangement seen in KSRP KH2 and KH3 modules, with the relative orientation stabilized mainly by domain-linker contacts (44). Comparison with the KH module arrangement observed in KHNYN and N4BP1 now defines a new group of interdomain interaction with an orthogonal packed conformation. This differs from the orthogonal linker group because it has a different absolute rotational orientation and the arrangement is stabilized by the packing between KH modules in the KHNYN and N4BP1 group instead of domain-linker interactions.

N4BP1 is an ISG (10) that has a similar domain arrangement to KHNYN (**Fig 2A**). Moreover, it has previously been shown to regulate ZAP antiviral activity in a genetic screen (45). However, it has not been shown to physically interact with ZAP and is predominantly localized to the nucleus (46). Given the structural similarity of the ex-diKH domains, we determined whether KHNYN and N4BP1 have redundant or complementary activity by siRNA knockdown of N4BP1 in CRISPR control or KHNYN CRISPR knockout cells in the absence (**Fig 2B)** and presence (**Fig 2C)** of type I interferon treatment. While there was no substantial increase in HIV-CpG infectious virus production when N4BP1 was knocked down in control cells, N4BP1 depletion increased HIV-CpG infectious virus to similar levels as HIV-WT in KHNYN CRISPR cells, regardless of interferon induction (**Fig 2B-C** and **Fig S2A-B**). This indicates that N4BP1 has a small amount of antiviral activity on HIV-CpG and there is some functional redundancy between the two proteins.

**Figure 2.**
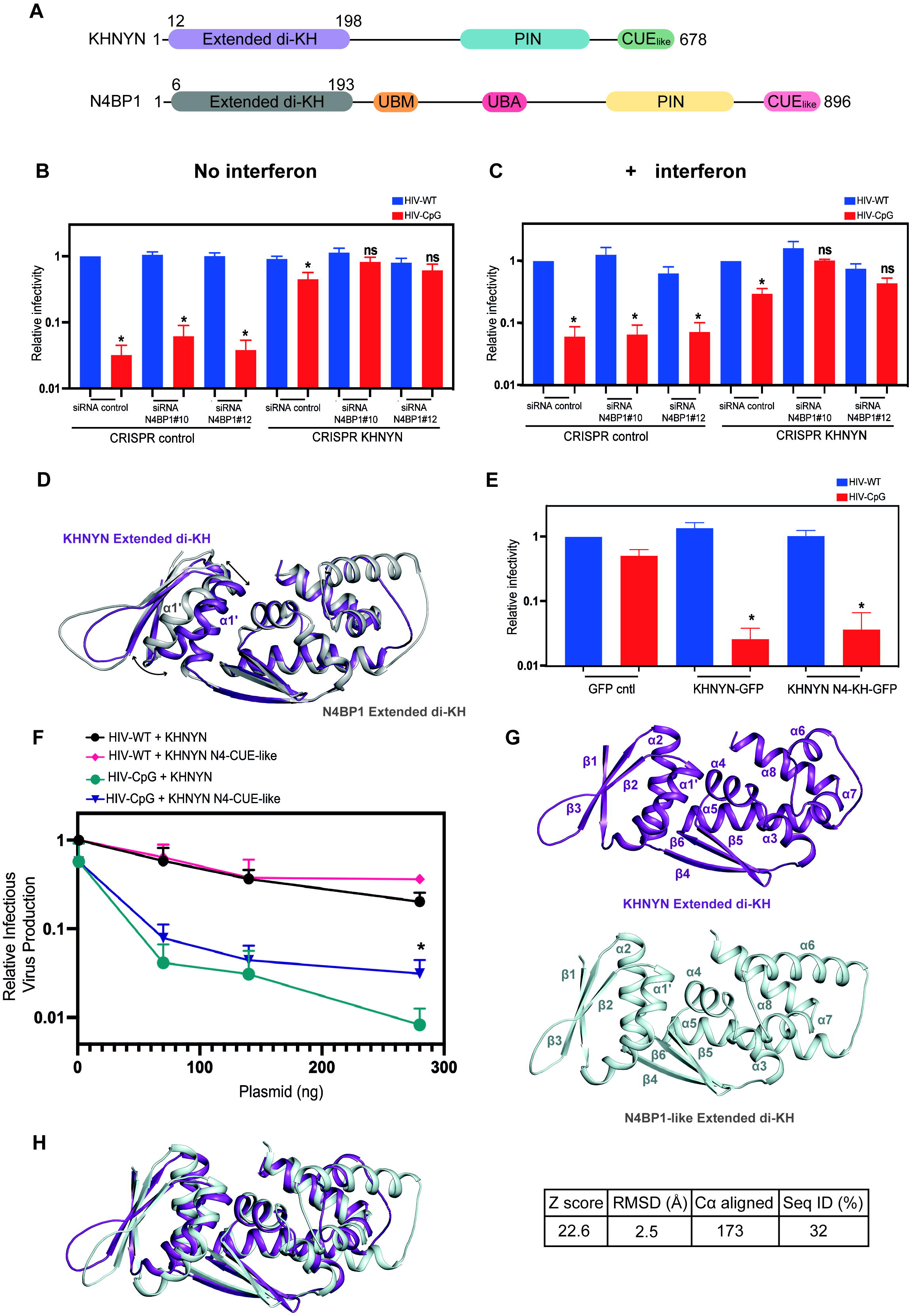
The KHNYN and N4BP1 ex-diKH domains are functionally equivalent. (**A**) Schematic representation of KHNYN and N4BP1. The position of structural/functional domains are highlighted; lilac (KHNYN ex-diKH domain residues 12-198), cyan (KHNYN PIN domain), green (KHNYN CUE_like_ domain), grey (N4BP1 ex-diKH domain residues 6-193), orange (N4BP1 UBM domain), deep pink (N4BP1 UBA domain), yellow (N4BP1 PIN domain) and pink (N4BP1 CUE_like_ domain). (**B-C**) Infectious virus production from HeLa CRISPR control or CRISPR KHNYN cells that were either untreated (**B**) or treated with 500 U of IFN-β (**C**) and transfected with either siRNA control or siRNAs against N4BP1 and infected with HIV-WT or HIV-CpG at MOI = 3. Each bar shows the average value of three independent experiments normalized to the value obtained for HIV-WT in the CRISPR control cells transfected with siRNA control. (**D**) 3D DALI structural superimposition of KHNYN and N4BP1 (PDB: 6Q3V) ex-diKH domains. Structures, shown in cartoon representation, KHNYN in purple and N4BP1 in grey, were aligned over all backbone Cα atoms (Z score 23.9, RSMD = 1.8 Å over 174 Cα) (**E**) Infectious virus production from stable HeLa CRISPR KHNYN GFP control, KHNYN WT-GFP or KHNYN N4-diKH-GFP infected with either HIV-1 WT or HIV-1 CpG at a MOI = 3. Bar plots represent the average value of three independent experiments normalized to the value obtained for HIV-WT in the absence of KHNYN. (**F**) Infectious virus production from KHNYN CRISPR HeLa cells co-transfected with pHIV-WT or pHIV-CpG and increasing amount of CRISPR-resistant flag-tagged pKHNYN or pKHNYN N4-CUE-like expressing plasmids. Each point shows the average value of three independent experiments normalized to the value obtained for HIV-WT in the absence of KHNYN. (**G**) Structure of KHNYN ex-diKH domain (top) and Alphafold model of the ex-diKH domain from *Branchiostoma belcheri* NEDD4-binding protein 1-like (NBP1-like) (bottom). Protein backbones are shown in cartoon, secondary structure elements are labelled from N- to C-termini. (**H**) 3D DALI structural superimposition of KHNYN and N4BP1-like ex-diKH domains. Protein backbones are shown in cartoon representation, KHNYN in purple and N4BP1-like in pale cyan. Structures were aligned over all backbone Cα atoms. Alignment data is shown on the right (Z score = 22.6, rmsd= 2.5 Å over 173 Cα, 32 % Seq ID)

Because KHNYN displays greater antiviral activity than N4BP1, even though it shares the same orthogonal packed diKH domain arrangement as well as the CTB extension, we examined the 3D superimposition of the two ex-diKH structures in detail. This analysis showed that the only substantial difference between the two structures is a small displacement (3-4 Å) in the backbone of the KH1 α1’ helix of KHNYN into the KH1-KH2 interface when compared to N4BP1 (**Fig 2D**). Therefore, to determine if there are functional differences between these domains, infectious virus production was measured in stable cell lines expressing wild type KHNYN-GFP (24) or expressing KHNYN with the ex-diKH domain replaced with that from N4BP1 (KHNYN N4KH-GFP). These data (**Fig 2E**) reveal no significant difference in infectious virus production, indicating that these domains are functionally equivalent. By comparison, our previous characterization of KHNYN had implicated the CUE_like_ domain to be required for KHNYN function and also to have diverged evolutionary from the N4BP1 CUE_like_ domain (24). Swapping the CUE_like_ domains significantly decreased KHNYN activity (**Fig 2F** and **Fig S2C**), suggesting that KHNYN is more antiviral than N4BP1 because the CUE_like_ domain contains a nuclear export signal present in the KHNYN tetrapod lineage and not in N4BP1 (24).

We then considered whether the ex-diKH is a recent evolutionary innovation or present in ancient N4BP1-like proteins that predate the gene duplication that led to divergent N4BP1 and KHNYN proteins in bony fish (24). Lancelets (Cephalochordata) are a basal chordate that diverged from other chordates, including vertebrates, over 500 million years ago (47) and encode one N4BP1-like protein (NCBI gene IDs: Branchiostoma belcheri 109480521; Branchiostoma floridae 118417762 and Branchiostoma lanceolatum 136438319). Inspection of the AlphaFold model for the Branchiostoma belcheri N4BP1-like protein (AF-A0A6P4ZNE3-F1-v4) showed that an ex-diKH domain was present with a similar overall structure to the KHNYN ex-diKH domain (**Fig 2G-H**). This indicates that the ex-diKH domain is an evolutionary ancient domain that is present in N4BP1-like proteins throughout the chordate lineage.

### The KHNYN extended-diKH domain has no detectable RNA binding activity

Next, we examined the KHNYN ex-diKH domain potential for RNA binding. In structures of nucleic acid bound KH complexes, typically the single stranded nucleic acid backbone binds the GxxG motif in an extended conformation, with the Watson-Crick edge of the nucleic acid oriented towards the central beta sheet of the KH domain (34, 48, 49). This positions four unpaired nucleic acid bases into a hydrophobic binding cleft where Van der Waals, electrostatic interactions and hydrogen bonding stabilize the binding of nucleic acid (34). KH1 does not contain the GxxG motif usually located between α1 and α2 in a KH module (**Fig 3A**). Instead, the two helices form a continuous helix α1’ connected by the proline kink link at P29, suggesting KH1 has lost the capacity to bind RNA. In KH2, an intact GxxG motif (residues 94-97, GAQG) is present and forms the canonical loop connecting the α1 and α2 helices (α3 and α4 in **Fig 1C**). To determine the functional relevance of the KH2 GxxG motif, it was mutated to GDDG, which has been shown to prevent RNA binding in other KH domains (48). Surprisingly, this mutation had no effect on KHNYN antiviral activity (**Fig 3B** and **Fig S3A**) or subcellular localization (**Fig S3B**), suggesting that KH2 RNA binding activity is not required for KHNYN antiviral activity or that KH2 does not bind RNA.

**Figure 3.**
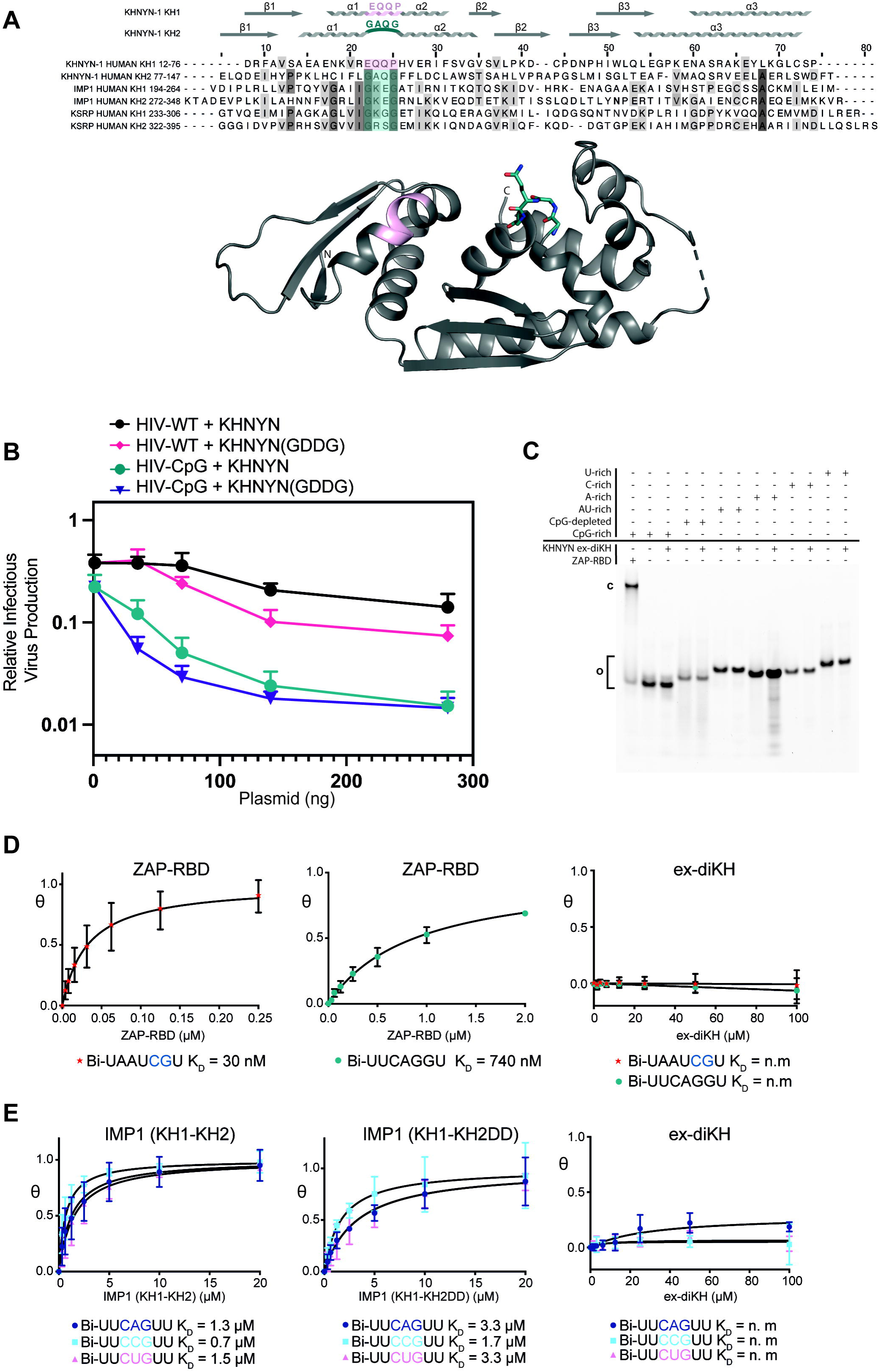
The KHNYN ex-diKH domain has no detectable RNA binding activity. (**A**) (Upper) Multiple sequence alignment of KH domains from human KHNYN (Uniprot: O15037), Insulin-like growth factor 2 mRNA binding protein 1 (IMP1) (Uniprot: Q9NZI8) and KH type-splicing regulatory protein (KSRP) (Uniprot Q92945). Conserved residues are highlighted with increasing shades of grey. GxxG motifs are highlighted in cyan and the equivalent residues in pink for KHNYN KH1. KHNYN KH1 and KH2 secondary structure elements aligned with the sequence are shown above. (Lower) KHNYN ex-diKH domain shown in cartoon representation. The position of the proline-kinked helix in KH1 and GxxG motif in KH2 are colored according to the scheme above with the GxxG motif (^94^GAQG^97^) also highlighted in sticks. (**B**) Infectious virus production from KHNYN CRISPR HeLa cells co-transfected with pHIV-WT or pHIV-CpG and increasing amount of CRISPR-resistant flag-tagged pKHNYN or pKHNYN(GDDG) expressing plasmids. Each point shows the average value of three independent experiments normalized to the value obtained for pHIV-WT in the absence of KHNYN. (**C**) EMSA analysis of ex-diKH RNA-binding. U-rich, C-rich, A-rich, AU-rich, CpG-depleted, and CpG-rich 33mer 5’-FAM-labeled RNAs (0.2 µM) were incubated with ZAP RBD (2 µM) and ex-diKH (20 µM). Free (**o**) and bound (**c**) components were separated on a 10% acrylamide native TBE gel and visualized by fluorescence imaging. (**D & E**) BLI analysis for (**D**) KHNYN ex-diKH and ZAP-RBD binding to CpG (Bi-UAAUCGU) and non-CpG (Bi-UUCAGGU) RNAs and (**E**) KHNYN ex-diKH, IMP1(KH1-KH2) and IMP1(KH1-KH2DD) RNA-binding to 5’-biotinylated CNG RNAs. Non-linear fits of fraction bound plotted against protein concentration are shown. Data are the mean of triplicate measurements. Error bars represent the standard deviation of the mean for each triplicate measurement. The derived *K_D_* values are shown beneath plots and presented with 95 % CI values in **Table S4**. BLI signal response for ex-diKH interaction with CpG, non-CpG and CNG RNAs is unobservable.

To differentiate between these two possibilities, an electrophoretic mobility shift assay (EMSA) was performed using 33mer 5’-FAM-labeled U-, C-, A-, AU-, CpG-rich and CpG-depleted RNAs (**Table S3**). As a control, ZAP RBD bound efficiently to the CpG-rich RNA. However, the KHNYN ex-diKH did not bind to any of the RNA sequences even at high excess of protein (100-fold) (**Fig 3C**). To further analyze RNA binding, comparative measurements with short RNAs (**Table S4**) were made using bio-layer interferometry (BLI) experiments with KHNYN ex-diKH, ZAP-RBD and the KH1-KH2 di-domain of IMP1 that scored as a top match in the DALI search (**Table S2**). These data showed that the ZAP RBD bound a short CpG containing RNA with high affinity (*K_D_* = 30 nM) and to an RNA without a CpG with a 25-fold lower affinity (*K_D_* = 740 nM) (**Fig 3D, Fig S3C,** and **Table S4**). By comparison, the KHNYN ex-diKH showed no measurable interaction with either RNA (**Fig 3D, Fig S3C** and **Table S4**). Further, BLI titrations with the IMP1 KH1-KH2 di-domain and a derivative containing a GDDG mutation in KH2 (43, 48) with CAG, CCG and CUG containing RNAs showed both IMP1 constructs bound these RNAs with low µM affinities with the KH2 GDDG mutation reducing the binding affinity around 2-fold (**Fig 3E, Fig S3C** and **Table S4**). Interaction between the KHNYN ex-diKH and RNA was again undetectable with any of these RNAs. Overall, no detectable RNA binding activity for the KHNYN ex-diKH domain was observed using a variety of RNA oligonucleotides employing orthogonal EMSA and BLI assays and at high protein excess. Although not every possible RNA sequence has been analyzed, these data indicate that the KHNYN ex-diKH domain does not bind RNA.

### The KHNYN extended-diKH domain may be a protein binding site

Given the lack of KHNYN ex-diKH domain RNA binding activity, an alternative hypothesis is that it is required for a protein-protein interaction. Further analysis of the inter-residue contacts in the ex-diKH crystal structure identified exposed arginine residues, R33 and R65, that make salt bridges and cation-π stacking interactions at the edge of the KH1-KH2 cleft (**Fig 4A**) that potentially could form part of a protein binding site. Mutation of these residues to abolish the interactions is likely to reduce the packing at the interdomain cleft, leading to uncoupling of KH1 and KH2 with potential functional consequences for the binding of protein interaction partners. Therefore, to test the requirement of these residues in KHNYN antiviral function, R33 and R65 were substituted to alanine either individually or in combination and the mutant proteins tested for antiviral activity (**Fig 4B-C**). While mutating R33 or R65 individually had no effect, when mutated in combination they substantially reduced KHNYN antiviral activity, even though the proteins were efficiently expressed (**Fig 4D**). These data suggest that the hydrogen bonding configuration of R33 and R65 and the exposed surface of the ex-diKH domain around the KH1-KH2 cleft is required for antiviral activity. Given the lack of RNA binding activity, this supports the notion that the ex-diKH domain serves as a protein-protein interaction domain with the KH1-KH2 cleft acting as a protein-binding site.

**Figure 4.**
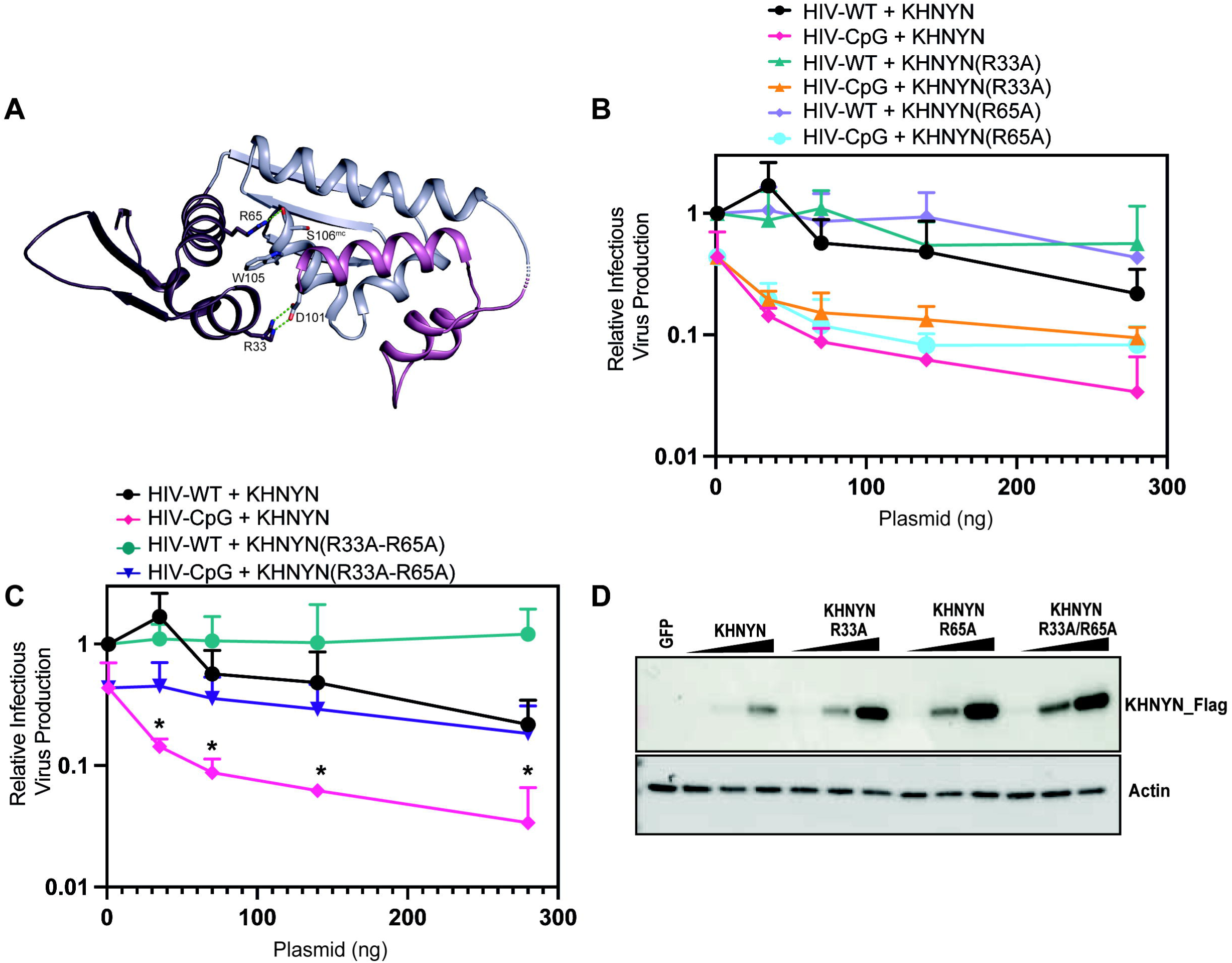
The KHNYN KH1-KH2 interdomain cleft is required for antiviral activity. (**A**) The KHNYN ex-diKH domain is shown in cartoon representation. KH1, KH2 and CTB are colored purple, blue grey and magenta respectively. Residues R33 and R65 together with D101, W105 and S106 that make salt bridge, hydrogen bond and cation-π interactions across the KH1-KH2 interdomain cleft are shown in sticks. Salt bridge and hydrogen bonds are represented by the green dashes, mc = mainchain. (**B-C**) Infectious virus production from KHNYN CRISPR HeLa cells co-transfected with pHIV-WT or pHIV-CpG and (**B**) increasing amount of CRISPR-resistant FLAG-tagged pKHNYN, pKHNYN(R33A) or pKHNYN(R65A) and (**C**) pKHNYN or pKHNYN(R33A/R65A) expressing plasmids. Each point shows the average value of three independent experiments normalized to the value obtained for pHIV-WT in the absence of KHNYN. (**D**) Representative western blotting corresponding to the expression level of FLAG-tagged KHNYN KHNYN(R33A), KHNYN(R65A) or KHNYN(R33A-R65A) employed in the infection experiments shown panels **B** and **C**.

## Discussion

In this study, we have structurally and functionally characterized the KHNYN ex-diKH domain. The structure identifies it as a member of a new group of type-1 diKH domains, the orthogonal packed, based on the relative orientation of the two KH modules. In addition, the KHNYN ex-diKH domain contains an extension to the KH modules comprising a C-terminal helical bundle that packs into KH2. The diKH arrangement and C-terminal extension represent a novel domain architecture observed only in KHNYN and its paralogs and is divergent from previously observed diKH arrangements. Although the KH modules of KHNYN adopt the classic secondary structure fold of type-I KH domains, KH1 lacks a GxxG motif, while it is retained in KH2. This elicits structural variation between the KH modules in KHNYN, with the GxxG loop missing in KH1 replaced with a proline kink, indicating further structural divergence from classical KH domains. Notably, we detect no RNA binding activity, suggesting that the orthogonal packed diKH family may have evolved to function outside of the usual KH RNA binding activity.

The function of the ex-diKH domain in the context of KHNYN, N4BP1 or NYNRIN is unknown. However, this domain is required for KHNYN antiviral activity. Our data shows that while both KH modules in the ex-diKH domain have a type-1 KH fold, they do not bind RNA in EMSA or BLI experiments. Moreover, this domain also does not regulate KHNYN subcellular localization or its interaction with ZAP and does not mediate homo-multimerization. Inspection of the crystal structure highlighted a potential protein binding site at the KH1-KH2 interface and mutations that disrupt salt bridges within this site substantially decreased KHNYN antiviral activity. Therefore, we hypothesize that the KHNYN ex-diKH domain may act as a protein-protein interaction unit that binds a currently unknown cofactor in the ZAP antiviral system. Identifying binding partners for KHNYN may identify novel cofactors for ZAP-mediated RNA decay. Furthermore, since the N4BP1 ex-diKH domain functionally substitutes for the KHNYN ex-diKH domain, such interactions are likely conserved and could represent an ancient function pre-dating the evolution of ZAP.

## Methods

### Plasmids and Cell lines

HeLa, HEK293T and TZM-bl cells were maintained in high glucose DMEM supplemented with GlutaMAX (Thermo Fisher Scientific), 10% fetal bovine serum, 100 U/mL penicillin and 100 µg/mL streptomycin and incubated with 5% CO2 at 37°C. Control CRISPR and KHNYN CRISPR HeLa cells were previously described (18). The CRISPR-resistant pKHNYN-FLAG plasmid has been previously described (18). Specific mutations were created using site-specific mutagenesis. The KHNYN deletion mutants were generated by site-directed mutagenesis. The KHNYN domain swap constructs were generated by HiFi assembly (NEB) using amplicons obtained using Q5 DNA polymerase (NEB). The generated plasmids were verified by Sanger sequencing. All primers are shown in **Table S5** and all PCR reactions were performed using Q5 High-Fidelity polymerase (NEB). HIV-1_NL4-3_ (pHIV-WT) and HIV_env86-561CpG_ (pHIV-CpG) in pGL4 were previously described (18, 50). Stable CRISPR KHNYN HeLa cells expressing KHNYNΔex-diKH-GFP and KHNYN-N4KH-GFP were produced by transduction with the murine leukemia virus (MLV) retroviral vector MIGR1 (51) with the KHNYN-GFP constructs cloned into the multiple cloning site and GFP replaced by the Blasticidin S-resistance gene.

### Transfections and infections

HeLa cells were seeded in 24-well plates at 70% confluency. Cells were transfected according to the manufacturer’s instructions using TransIT-LT2020 (Mirus) at the ratio of 3 µL TransIT-LT2020 to 1 µg DNA. 0.5 µg pHIV-WT or pHIV-CpG and the designated amount of KHNYN-FLAG or GFP-FLAG for a total of 1 µg DNA were transfected. 24 hours post-transfection, the culture media was replaced with fresh media. Supernatants and cells were harvested for analysis 24 hours later. For infection experiments, viral stocks were produced by co-transfecting pHIV-WT or pHIV-CpG with pVSV-G (52) into HEK293T ZAP CRISPR cells (53) and titered on TZM-bl cells (54–56). HeLa cells were plated as for transfection experiments and infected either with HIV-WT or HIV-CpG at MOI = 3. Media was replaced 24 hours after infection. Supernatants and cells were harvested 48 hours post-infection.

### TZM-bl infectivity assay

The TZM-bl indicator cell line was used to quantify the amount of infectious virus (54–56). Briefly, cells were seeded in 96-well plates and infected by incubation with virus stocks or supernatants of infected cells. 48 hours post-infection, the cells were lysed and infectivity was measured by β-galactosidase expression using the Galacto-Star System following manufacturer’s instructions (Applied Biosystems). β-galactosidase activity was quantified as relative light units per second using a PerkinElmer Luminometer.

### Analysis of protein expression by immunoblotting

48 hours post-transfection, HeLa cells were lysed in Laemmli buffer at 95°C for 10 minutes. The culture supernatant was also collected, and virions pelleted by centrifugation for 2 hours at 20,000 x g through a 20% sucrose cushion in phosphate-buffered saline (PBS). Viral pellets were resuspended in 2x Laemmli buffer. Cell and virion lysates were resolved on 8 to 16 % Mini-Protean TGX precast gels (Bio-Rad), transferred into nitrocellulose membranes (GE Healthcare) that were then blocked in 5 % non-fat milk in PBS supplemented with 0.1% Tween 20. Primary antibodies were incubated overnight at 4°C followed by three washes with PBS and then corresponding secondary antibody (1:5000 anti-mouse IRDye 680RD (LI-COR, 926–68070), 1:5000 anti-rabbit IRDye 800CW (LI-COR, 926–32211), 1:5000 anti-rabbit HRP (Cell Signaling Technology, 7074) or 1:5000 anti-mouse HRP (Cell Signaling Technology, 7076)) was incubated for one hour. Proteins were visualized by LI-COR (Odyssey Fc) measuring secondary antibody fluorescence or using Amersham ECL Prime Western Blotting Detection reagent for HRP-linked antibodies with an ImageQuant (LAS8000 Mini). The following primary antibodies were used at dilutions of: 1:50 HIV anti-p24Gag (mouse hybridoma) (57), 1:3000 anti-HIV gp160/120 (Rabbit, ADP421; Centralized Facility for AIDS Reagents (CFAR), 1:5000 anti-HSP90 (Rabbit, GeneTex, GTX109753), 1:1000 anti-FLAG (DYKDDDDK, Rabbit, Cell Signaling, 14793), 1:5000 anti-ZAP (Rabbit, Abcam, ab154680), 1:2000 anti-β-actin (Mouse, Abcam; ab6276), 1:10000 anti-GAPDH (Rabbit, Proteintech, 10494-1-AP), 1:10000 anti-GAPDH (mouse, Proteintech, 60004-1-Ig), 1:1000 anti-GFP (Mouse, Roche 11814460001).

### Immunoprecipitation assays

HeLa cells stably expressing wild type KHNYN-GFP or mutant proteins were seeded in 6-well plates 24 hours prior to immunoprecipitation. The cells were lysed on ice in 500 µL of lysis buffer (0.5% NP-40, 150 mM KCl, 10 mM HEPES pH 7.5, 3 mM MgCl_2_) supplemented with complete Protease inhibitor cocktail tablets (Roche 11873580001). The lysates were incubated on ice for 1 hour and centrifugated at 20,000 x g for 15 minutes at 4°C to pellet the nuclear fraction. 50 µL of the post-nuclear supernatants was saved as the input lysate and 450 µL were incubated with 5 µg of anti-GFP antibody (Roche 11814460001) for 1 hour at 4°C. Protein G Dynabeads (Invitrogen) were then added and incubated overnight at 4°C with rotation. The lysates were washed four times with wash buffer (0.05% NP-40, 150 mM KCl, 10 mM HEPES pH 7.5, 3 mM MgCl_2_) before the bound proteins were eluted with 2x Laemmli buffer and boiled for 10 minutes. Protein expression was analyzed by western blotting as described above.

### Microscopy

HeLa cells were seeded in 24-well plates on coverslips pre-treated with poly-lysine. KHNYN CRISPR HeLa cells were transfected with 250 ng of KHNYN-FLAG or KHNYN(GDDG)-FLAG. 24 hours post-transfection, the cells were fixed with 4% paraformaldehyde for 20 minutes at room temperature, washed once with 1xPBS and once in 10 mM glycine. Cells were then permeabilized for 15 minutes in 1% BSA and 0.1% Triton-X in PBS. HeLa cells stably expressing wild type KHNYN-GFP or versions with specific mutations were seeded in pre-treated 24-well plates 24 hours prior to immunostaining and fixed and permeabilized as above. Mouse anti-FLAG (1:500), rabbit anti-ZAP (1:500) or rabbit anti-TRIM25 (1:500) antibodies were diluted in PBS/0.01% Triton-X and the cells were stained for 2 hours at room temperature. The cells were then washed three times in PBS/0.01% Triton-X and incubated with Alexa Fluor 594 anti-mouse or anti-rabbit or Alexa Fluor 647 anti-rabbit (Molecular Probes, 1:500 in PBS/0.01% Triton-X) for 45 minutes in the dark. Finally, the coverslips were washed three times with 1xPBS/0.01% Triton-X and then mounted on slides using Prolong Diamond Antifade mountant with DAPI (Invitrogen). Imaging was performed on a Nikon Eclipse Ti Inverted Microscope, equipped with a Yokogawa CSU/X1-spinning disk unit, under 60x objective and laser wavelengths of 405 nm, 488 nm, 561 nm, and 640 nm. Image processing and co-localization analysis was performed with Image J (Fiji) software.

### Leptomycin B treatment

HeLa cells stably expressing wild-type KHNYN-GFP or mutant versions were seeded in pre-treated 24-well plates 24 hours prior to 4-hour treatment with 50 nM of Leptomycin B or DMSO at 37°C. After treatment, the cells were fixed and immuno-stained as described above.

### Protein expression and purification

DNA sequences encoding human KHNYN residues P8-G200 and human ZAP residues M1-N227, were PCR amplified and inserted into a pET-52b(+) expression vector (Novagen), using XmaI/SacI restriction sites (KHNYN) or ligation independent cloning (ZAP), to produce N-terminally StrepII-tagged fusion proteins. Primer sequences are listed in **Table S5**. Insert sequences were confirmed by DNA sequencing (Eurofins). StrepII-tagged fusion proteins were expressed in *E. coli* strain BL21(DE3) (NEB) incubated at 37°C with shaking. Addition of 0.4 mM IPTG was used to induce protein expression at an A_600_ of ∼0.6, followed by 18 hour incubation at 18°C. Cells were harvested by centrifugation, resuspended in 5 mL/g of lysis buffer K (KHNYN(8-200)) or lysis buffer Z (ZAP(1-227)) and lysed by sonication (KHNYN(8-200)) or Emulsiflex homogenization (ZAP(1-227)). The composition of protein purification buffers is presented in **Table S6**. Lysate was clarified by centrifugation for 1 hour at 50,000 xg at 4°C before the supernatant was applied to 10 mL of packed Strep-Tactin XT Superflow resin (IBA), followed by washing with 50 mL of wash K1 buffer followed by 50 mL wash K2 buffer (KHNYN-1 (8-200)) or 200 mL of wash Z1 buffer (ZAP(1-227)) at 4°C. Bound protein was then step eluted with elution buffer KE (KHNYN(8-200)) or ZE (ZAP(1-227)) and highest purity fractions, identified by SDS PAGE, pooled.

For KHNYN(8-200), GST-tagged HRV 3C-protease (0.5 mg) was added to eluent overnight to remove the StrepII-tag. Following concentration of cleaved eluent, GST-tagged HRV 3C-protease was removed by affinity chromatography using a 1 mL GSTrap HP column (Cytiva) before sample application to a Superdex 75 26/600 column (Cytiva) equilibrated in KGF gel filtration buffer. Peak fractions were identified by SDS PAGE, concentrated to ∼30 mg/mL and flash frozen in liquid nitrogen.

For ZAP(1-227), GST-tagged HRV 3C-protease (0.5 mg) was added to the eluent and that was then dialysed overnight against Low Salt buffer. The dialysate was then diluted rapidly to 125 mM NaCl before application to a 1 mL HiTrap Q HP column (GE). The flow through was collected along with 10 mL of wash with IEX buffer. This sample was concentrated and any remaining GST-tagged HRV 3C-protease was removed by affinity chromatography using a 1 mL GSTrap HP column (GE) before sample application to a Superdex 75 16/600 column (GE) equilibrated in ZGF gel filtration buffer. Peak fractions identified by SDS PAGE were then concentrated to ∼2.5 mg/mL and flash frozen in liquid nitrogen. IMP1 proteins were purified as described (43) and kindly provided by Dr Robert Dagil, University of Copenhagen.

### SEC-MALLS

Size exclusion chromatography coupled to Multi-Angle Laser Light Scattering (SEC-MALLS) was performed to determine the molar mass of KHNYN(8-200) samples. 100 µL samples ranging from 0.5 to 8mg/ml were applied to a Superdex 200 10/300 INCREASE GL column (Cytiva) equilibrated in 50 mM BTP pH 8.5, 150 mM NaCl, 0.5 mM TCEP and 3DmM NaN_3_ at a flow rate of 1.0 mL/min. The scattered light intensity and protein concentration of the column eluate were recorded using a DAWN-HELEOS-II laser photometer and an OPTILAB-TrEX differential refractometer (dRI) (*dn/dc* = 0.186) respectively. The weight-averaged molecular mass of the protein was then determined using the ASTRA software version 7.3.2 (Wyatt Technology Corp.).

### Crystallization and structure determination

KHNYN (8-200) crystals were produced by vapour drop diffusion at 18°C using protein at stock concentration (∼30 mg/mL) in Swissci MRC two-drop 96-well plates (Molecular Dimensions), with a 75 µL reservoir of screen condition (100 mM Tris pH 7.0, 250 mM NaCl and 35% w/v PEG 3K). A Mosquito LCP robot (TTP Labtech) was utilized to set 100 nL drops, with a 2:1 ratio of protein solution to screen condition. Crystals were harvested by looping and flash-frozen in liquid nitrogen before data collection on the i04 beamline at Diamond Light Source at a wavelength of 0.9795 Å.

An initial dataset was processed using the automatic xia2 dials Diamond pipeline (58, 59) and the structure solved by molecular replacement using PHASER (60) employing an AlphaFold2 model (35) of KHNYN residues 12-142. After initial rebuilding in COOT 0.9.6 (61) and iterative rounds of refinement in PHENIX REFINE (62), a higher quality and more complete dataset was obtained and processed with the autoPROC pipeline (63) using XDS (64), POINTLESS (65) and AIMLESS (66) and anisotropic scaling applied using STARANISO (http://staraniso.globalphasing.org/cgibin/staraniso.cgi) and inheriting the FreeR flags from the initial dataset. The final model comprises two copies of KHNYN(8-200) with all residues built from 12-198 except for a flexible loop between residues 148-159 and was refined to a R_work_/R_free_ of 0.226/0.282. MolProbity (67) and PDB_REDO (68) were used to monitor and assess model geometry throughout the refinement process. Details of crystal parameters, data collection and refinement statistics are shown in **Table S1**. The co-ordinates and structure factors have been deposited in the PDB under the accession code 9HTS.

### Structural and sequence alignments

Protein domain prediction was performed with AlphaFold2 (35) and Phyre2 (37). PDB files from crystal structures were viewed and rendered in PyMOL 2.5.5 (Schrodinger, LLC). Electron density maps were visualized in COOT 0.9.6 (61). Analysis of protein interfaces within crystal structures was performed with PDB-PISA (38). The DALI server (39) was used to perform PDB25/PDB90 searches using chain A of the KHNYN(8-200) crystal structure. Primary sequences were aligned in Jalview (69) using Clustal default settings. Secondary structure was annotated based on the KHNYN(8-200) crystal structure.

### siRNA transfections and IFN treatment

OnTarget Plus siRNA control and siRNA against N4BP1 were purchased from Horizon Discovery Biosciences Limited. HeLa CRISPR control or CRISPR KHNYN cells were reverse-transfected with either siRNA control or siN4BP1 as follows: 1 uL of each siRNA at 20 nM with 1 µL of Lipoafectamine RNA-iMax following manufacturing instructions. 24 hours after the first transfection, the media was replaced, and cells were transfected a second time, directly, using the same conditions. 48 hours after the first transfection, cells were infected with either HIV-WT or HIV-CpG at MOI 3. Media was replaced 24 hours post-infection. Supernatants and cells were then harvested 48 hours post-infection and analyzed by TZM-bl assay and by immunoblotting. For the IFN-β experiments, cells were treated with 500 U of IFN-β at 24 hours post-infection and harvested 24 hours later.

### RNA

HPLC purified RNA oligonucleotides were synthesized by Horizon Discovery Biosciences Limited. RNA was deprotected according to manufacturer’s protocol, lyophilized and then resuspended in TE buffer (10 mM Tris pH 8.0 and 0.1 mM EDTA). Oligonucleotide concentrations were determined from the A_260_ nm derived from the UV/Vis absorbance spectrum using extinction coefficients based on base sequence composition provided by the manufacturer.

### Electrophoretic mobility shift assays (EMSA)

To test ex-diKH RNA-binding, 10 µL reactions containing 0.2 µM of 33-mer 5’-FAM-labeled RNAs (**Table S3**) were incubated with 20 µM KHNYN(8-200) in EMSA reaction buffer (20 mM Tris pH 8.0, 50 mM NaCl, 0.5 mM TCEP, 5% w/v glycerol) together with a positive control sample containing 0.2 µM 5’-FAM-labeled CpG-rich RNA and 2 µM ZAP (1-227). Reactions were incubated at room temperature for 30 minutes before loading onto a 10% acrylamide TBE gel prior to electrophoresis for 1 hour in 1 X TBE running buffer (pH 8.3) at 30 mA and 125 V on ice. Free and bound nucleic acids were imaged using a Typhoon™ 9500 imager with the FAM-default sample detection method (λ_laser_ = 473 nm, λ_emmision_ = 520 nm, filter LPB) at gain between 600-700 V. Quantification of integrated band intensities was performed within the ImageQuant TL Software.

### Biolayer interferometry (BLI)

BLI was performed on an Octet R8 (Sartorius) instrument at 25°C. 10 nM of a panel of biotinylated RNAs (**Table S4**) in BLI buffer (20 mM Tris pH 8.0, 50 mM NaCl, 0.5 mM TCEP, 0.01% Tween 20 and 8 units/mL RNAsin Plus (Promega)) was immobilized onto Streptavidin-coated biosensors. RNA-bound biosensors were then incubated with varying protein concentrations: 0-100 µM of KHNYN(8-200), 0-20 µM of IMP1(V194-N369) or IMP1(V104-N369, K294D, E295D) and either 0-0.25 µM or 0-2 µM of ZAP(1-227). Between reads with biosensors, three-step regeneration was performed with BLI buffer containing 1 M NaCl. BLI interferograms were processed in Octet BLI Analysis 12.2.2.4 software (Sartorius), double referencing data with 0 µM protein incubations and unloaded biosensors. Average responses were then calculated from the end point of association, converted to fraction bound (θ) based on the maximal response at saturation (R_max_) and plotted as a function of protein concentration. *K_D_* values were determined in GraphPad PRISM10 (GraphPad Software Inc.) by fitting to a one site specific hyperbolic binding curve (θ = B_max_*x/(*K_D_* + x)) where B_max_ is the maximal response.

### Statistical analysis

For restriction assays, statistical significance was determined using unpaired two-tailed t test in GraphPad Prism10 (GraphPad Software Inc.). Data are represented as mean ± standard deviation and significance was ascribed to p values p < 0.05.

## Acknowledgements

We thank members of the Neil, Taylor and Swanson laboratories for helpful discussions and Dr Robert Dagil, University of Copenhagen for the gift of IMP1 proteins. We gratefully acknowledge the Francis Crick Structural Biology Science Technology platform and Diamond Light Source, Didcot, UK (Block allocation group MX25587) and beamlines i04 for access and support from Beamline staff. The following reagents were obtained through the NIH AIDS Research and Reference Reagent Program, Division of AIDS, NIAID, NIH: TZM-bl from Dr. John C. Kappes, Dr. Xiaoyun Wu and Tranzyme Inc; HIV-1 p24 Hybridoma (183-H12-5C) from Dr. Bruce Chesebro. The Antiserum to HIV-1 gp120 #20 (ARP421) was obtained from the NIBSC Centre for AIDS Reagents.

