## Supplementary data figures for "Structural and functional characterization of the KHNYN extended-diKH domain for mediating ZAP antiviral activity"

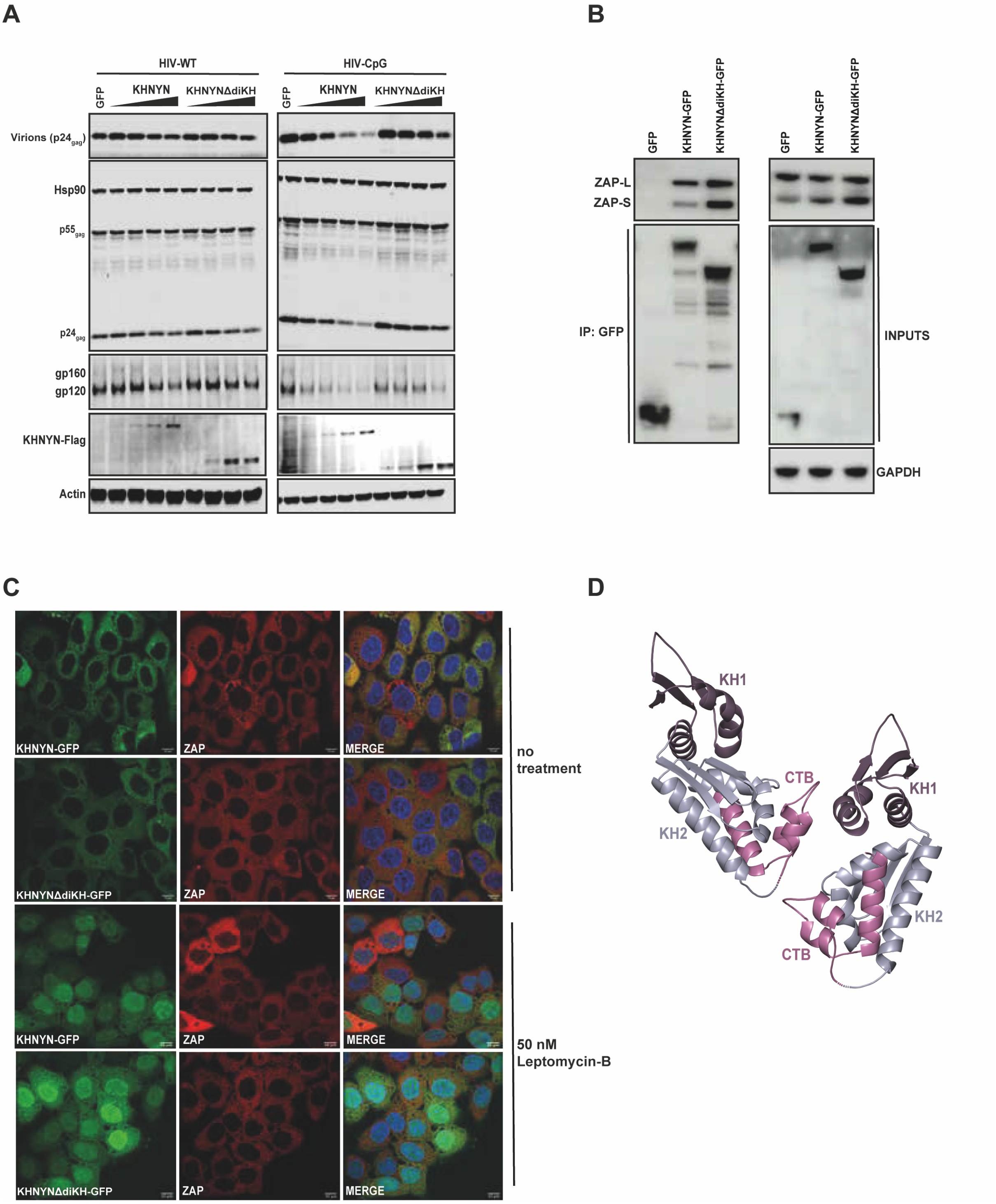


**Figure S1. The KHNYN ex-diKH domain is required for antiviral activity but does not regulate the interaction with ZAP or KHNYN subcellular localization.** (**A**) Representative western blotting corresponding to **Fig 1B**. (**B**) Representative ZAP-KHNYN co-immunoprecipitation experiment. Stable HeLa CRISPR KHNYN cells expressing GFP control, KHNYN-GFP or KHNYNΔdiKH-GFP were infected with either HIV-WT or HIV-CpG for 48 hours with MOI = 3. After infection, cells were lysed and immunoprecipitated using α-GFP antibodies. Samples were immunoblotted against GFP and ZAP. (**C**) Confocal microscopy staining for GFP and ZAP in HeLa CRISPR KHNYN cells expressing KHNYN-

GFP or KHNYNΔdiKH-GFP with no treatment or 50 nM of Leptomycin B for 4 hours at 37 °C. (**D**) The ex-diKH crystal structure asymmetric unit (ASU) relating to **Fig 1C**. The two ex-diKH protomers in the ASU are shown in cartoon representation. KH1, KH2 and CTB domains are colored purple, blue-grey and magenta respectively.

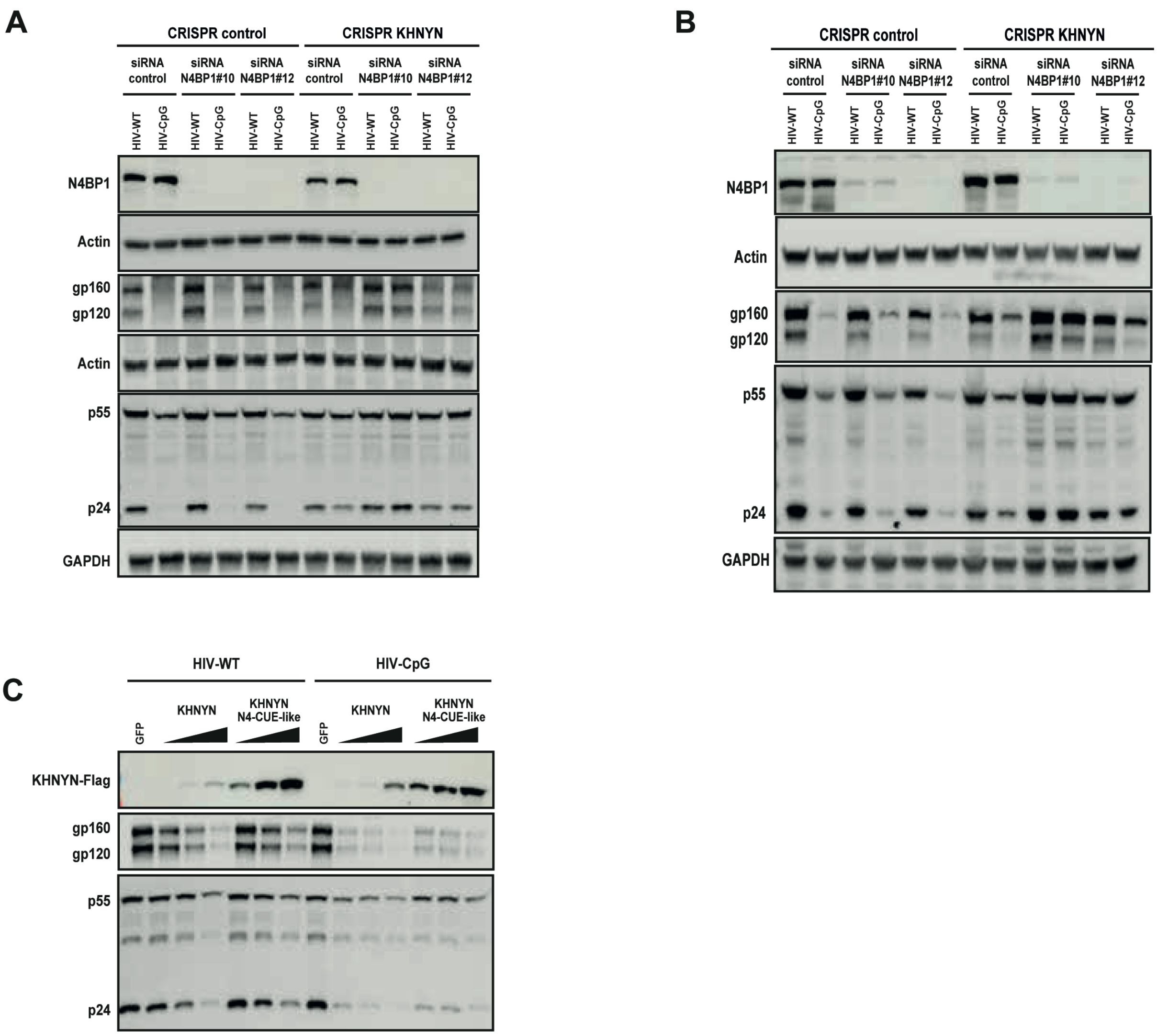


**Figure S2. The KHNYN and N4BP1 ex-diKH domains are functionally equivalent.** (**A**) Representative western blotting corresponding to **Fig 2B**. (**B**) Representative western blotting corresponding to **Fig 2C**. (**C**) Representative western blotting corresponding to **Fig 2F**.

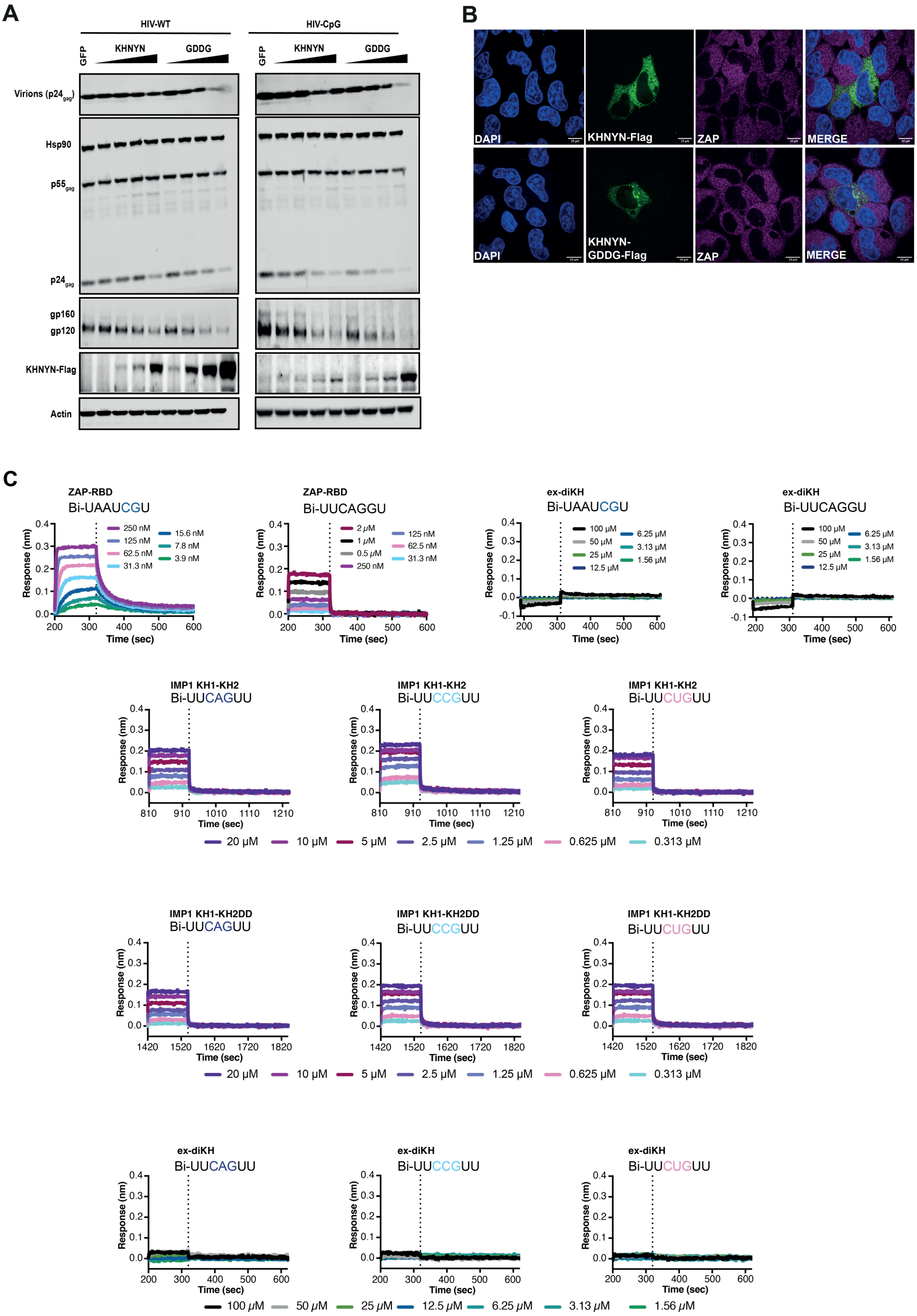


**Figure S3. The KHNYN ex-diKH domain has no detectable RNA binding activity.** (**A**) Representative western blotting corresponding to **Fig 3B**. (**B**) Confocal microscopy of

HeLa CRISPR-KHNYN expressing either FLAG-tagged KHNYN or KHNYN(GDDG) mutant. Cells were stained for the FLAG-tag (green) and ZAP (magenta). (**C**) BLI analysis of ZAP RBD, KHNYN ex-diKH, IMP1(KH1-KH) and IMP1(KH1-KH2DD) RNA binding, relating to **Fig 3D-3E**. BLI association and dissociation response traces are shown from a single titration from triplicate BLI runs performed with immobilized biotinylated RNAs and varying concentration of proteins. Switchover from the association to the dissociation phase is indicated by the dashed line.
