## Supplementary data tables for "Structural and functional characterization of the KHNYN extended-diKH domain for mediating ZAP antiviral activity"

#### Table S1. X-ray data collection and structure refinement statistics

| | KHNYN-1(P8-G200) Dataset 1 | KHNYN-1(P8-G200) Dataset 2 (STARANISO) |
|---|---|---|
| **Data collection** | | |
| Space group | $P2_1$ | $P2_1$ |
| Cell dimensions | | |
| a, b, c (Å) | 28.87, 122.64, 44.57 | 28.83, 121.80, 44.55 |
| α, β, γ | 90, 108.90, 90 | 90, 108.55, 90 |
| Wavelength (Å) | 0.9795 | 0.9795 |
| Resolution (Å) | 61.32-2.18 (2.22-2.18) | 60.90-2.20 (2.34-2.20) |
| Anisotropic diff. limits | | |
| a*, b*, c* (Å) | - | 2.68, 2.35, 2.04 |
| Unique reflections | 8927 (17) | 10863 (536) |
| $R_{meas}$ | 0.066 (0.280) | 0.188 (1.008) |
| $R_{pim}$ | 0.035 (0.198) | 0.079 (0.448) |
| $CC_{1/2}$ | 0.996 (1.000) | 0.995 (0.577) |
| I/σ(I) | 24.3 (2.2) | 7.5 (1.7) |
| Completeness | | |
| Spherical (%) | 58.7 (2.2) | 73.3 (23.0) |
| Ellipsoidal (%) | - | 90.8 (80.0) |
| Multiplicity | 3.3 (1.1) | 5.4 (4.8) |
| **Refinement** | | |
| Resolution (Å) | | 42.2 - 2.2 |
| $R_{work}/R_{free}$/Test set | | 0.226/0.282/0.072 |
| No. monomers/A.S.U. | | 2 |
| No. atoms | | |
| Protein | | 2764 |
| Water | | 28 |
| *B*-factors (Å) | | |
| Wilson | | 27.22 |
| Protein | | 28.11 |
| Water | | 25.46 |
| Average | | 28.08 |
| R.m.s. deviations | | |
| Bond lengths (Å) | | 0.005 |
| Bond angles (°) | | 0.97 |
| Chiral volumes ($Å^3$) | | 0.049 |
| Ramachandran | | |
| Favoured | | 96.52 |
| Allowed | | 3.19 |
| Outliers | | 0.29 |
| PDB code | | 9HTS |

†Values in parentheses refer to the highest resolution shell

**Table S2. KHNYN diKH DALI search results**

| DALI results, Z score >8.0 | Description | Z score | RMSD (Å) | Cα aligned | Seq ID (%) |
|---|---|---|---|---|---|
| 6q3v | N4BP1 KH1KH2 | 23.9 | 1.8 | 174 | 37 |
| 6qey | IMP1 KH1KH2 | 9.6 | 5.9 | 69 | 13 |
| 2n8l | IMP1 KH3KH4(DD) | 9.3 | 4.0 | 71 | 15 |
| 1ec6 | Nova-2 KH3 | 9.0 | 3.0 | 68 | 6 |
| 2jvz | KSRP KH2KH3 | 8.9 | 2.4 | 66 | 12 |
| 5www | MEX3C KH1 | 8.9 | 2.9 | 79 | 16 |
| 4aim | PNPASE KH | 8.8 | 9.2 | 76 | 5 |
| 2qnd | FMRP KH1KH2 | 8.3 | 1.7 | 63 | 16 |
| 2hh2 | KSRP KH4 | 8.2 | 2.7 | 65 | 12 |
| 1tua | APE0754 KH | 8.1 | 4.0 | 79 | 16 |
| 5wwx | MEX3C KH2 | 8.1 | 2.0 | 60 | 15 |

**Table S3. EMSA RNA oligonucleotides**

| Name | Sequence |
|---|---|
| CpG-rich | FAM-5'-GCAACGACGACGCUAUUUUGCGCAUCAGACGCG-3' |
| CpG-depleted | FAM-5'-GCAACCACCACUCUAUUUUGUGCAUCAGAUGCU-3' |
| AU-rich | FAM-5'-GUAACUAAUACUCUAUUUUGUUAAUCAGAUAAU-3' |
| A rich | FAM-5'-AUAACAAAUACACUAAAAUACAAAUCAGAAAAU-3' |
| C-rich | FAM-5'-CUCCUCCCUACACUCCCCUCUCCCGCCGCCCCU-3' |
| U-rich | FAM-5'-GUUUCUUUUACUCUUUGUUACUUUUCAGUUAAU-3' |

**Supplementary Table S4. BLI RNA binding experiments**

| Protein | Sequence | $K_D$ (μM) and 95 % CI |
|---|---|---|
| ZAP(1-227) | Bi-5'-UAAUCGU | 0.03 (0.02-0.06) |
| ZAP(1-227) | Bi-5'-UUCAGGU | 0.74 (0.54 – 1.01) |
| KHNYN-1 (8-200) | Bi-5'-UAAUCGU | n. m. |
| KHNYN-1 (8-200) | Bi-5'-UUCAGGU | n. m. |
| IMP1 (V194-N369) | Bi-5'-UUCAGUU | 1.3 (0.7 – 2.2) |
| IMP1 (V194-N369 | Bi-5'-UUCCGUU | 0.7 (0.5 – 1.0) |
| IMP1 (V194-N369 | Bi-5'-UUCUGUU | 1.5 (0.9 – 2.4) |
| IMP1 (V104-N369, K294D, E295D) | Bi-5'-UUCAGUU | 3.3 (2.0 – 5.4) |
| IMP1 (V104-N369, K294D, E295D) | Bi-5'-UUCCGUU | 1.7 (0.9 – 3.0) |
| IMP1 (V104-N369, K294D, E295D) | Bi-5'-UUCUGUU | 3.3 (1.9 – 5.6) |
| KHNYN-1 (8-200) | Bi-5'-UUCAGUU | n. m. |
| KHNYN-1 (8-200) | Bi-5'-UUCCGUU | n. m. |
| KHNYN-1 (8-200) | Bi-5'-UUCUGUU | n. m. |

### Table S5. Cloning primers

| Construct | Orientation | Sequence (5'-3') |
|---|---|---|
| KHNYN(8-200) | FWD | CAGTACGACCCGGGGCAAGTCCGGATCGTTTTGC |
| KHNYN(8-200) | REV | GCTAGACTGAGCTCTCATCAACCTGAGCTTGCTTCTTGC |
| ZAP(1-227) | FWD | CAGGGACCCGGTATGGCAGATCCGGAAGTTTGTTG |
| ZAP(1-227) | REV | GGCACCAGAGCGTTAATTTTTCTGCATGTGTTTGCTGTTG |
| KHNYNΔdiKH | FWD | GAGGCGTCTAGTGGGCAG |
| KHNYNΔdiKH | REV | CATGGTGAATTCGGTAGATCTAATTCC |
| KHNYN N4-KH-GFP | FWD | CCGAGCTCGGATCCGCCACCATGGCGGCCCGGGCGGTGCTG |
| KHNYN N4-KH-GFP | REV | GGTGGCGGATCCGAGCTC |
| KHNYN N4-CUE-like | FWD | ATGGTGTTGGAT GATTACAAGGATGACGACGATAAG |
| KHNYN N4-CUE-like | REV | TCTCTGAGCTGG ACTACCTTTTTCCTCTTCTCTCC |
| KHNYN N4-CUE-like | FWD | GAAAAAGGTAGT CCAGCTCAGAGATCTTCTGCAG |
| KHNYN N4-CUE-like | REV | ATCCTTGTAATC ATCCAACACCATGGCAGAAAG |
| KHNYN(GDDG) | FWD | CAAACTGCACTGCATCTTTCTGGGAGACGACGGCTTCTTCCTTGACTGCCTGGCCTG |
| KHNYN(GDDG) | REV | 5’CAGGCCAGGCAGTCAAGGAAGAAGCCGTCGTCTCCCAGAAAGATGCAGTGCAGTTTG |
| KHNYN(R33A) | FWD | CCATGTGGAGGCCATCTTCAGCGTGGG |
| KHNYN(R33A) | REV | GGCTGCTGTTCCCGAACC |
| KHNYN(R65A) | FWD | AAACGCCAGCGCAGCCAAGGAGTACCTG |
| KHNYN(R65A) | REV | TCCTTGGGGCCCTCCAGC |

#### Table S6. Protein purification buffers

| Protein | Buffer Name | Buffer Composition |
|---|---|---|
| KHNYN(8-200) | Lysis Buffer K | 100 mM BTP-HCl pH 8.5, 150 mM NaCl, 0.5 mM TCEP, 5% w/v glycerol, 0.2% v/v Triton X-100, 1x cOmplete EDTA-free protease inhibitor cocktail tablet per 50 mL (Roche) and 0.4 units/mL benzonase nuclease (Millipore) |
| ZAP(1-227) | Lysis Buffer Z | 100 mM BTP-HCl pH 6.5, 800 mM NaCl, 0.5 mM TCEP, 5% w/v glycerol, 10 mM $MgCl_2$, 1x cOmplete EDTA-free protease inhibitor cocktail tablet per 50 mL (Roche), 2.5 units/mL universal nuclease (Pierce) and 1.7 units/mL salt active nuclease (Sigma) |
| KHNYN(8-200) | Wash K1 | 100 mM BTP-HCl pH 8.5, 150 mM NaCl, 0.5 mM TCEP |
| KHNYN(8-200) | Wash K2 | 100 mM BTP-HCl pH 8.5, 1 M NaCl, 0.5 mM TCEP |
| ZAP(1-227) | Wash Z1 | 100 mM BTP-HCl pH 6.5, 800 mM NaCl, 0.5 mM TCEP, 5% w/v glycerol |
| KHNYN(8-200) | Elution buffer KE | 100 mM BTP-HCl pH 8.5, 150 mM NaCl, 0.5 mM TCEP, 50 mM biotin |
| ZAP(1-227) | Elution buffer ZE | 100 mM BTP-HCl pH 6.5, 800 mM NaCl, 0.5 mM TCEP, 5% w/v glycerol, 50 mM biotin |
| KHNYN(8-200) | Gel filtration KGF | 50 mM BTP-HCl pH 8.5, 150 mM NaCl, 0.5 mM TCEP |
| ZAP(1-227) | Low Salt | 100 mM BTP-HCl pH 6.5, 250 mM NaCl, 0.5 mM TCEP, 5% w/v glycerol |
| ZAP(1-227) | IEX | 100 mM BTP-HCl pH 6.5, 125 mM NaCl, 0.5 mM TCEP, 5% w/v glycerol |
| ZAP(1-227) | Gel filtration ZGF | 50 mM BTP-HCl pH 6.5, 200 mM NaCl, 0.5 mM TCEP, 5% w/v glycerol |
